# Mitochondrial position responds to glucose stimulation in a model of the pancreatic beta cell

**DOI:** 10.1101/2025.02.13.637960

**Authors:** Luis Perez, Xue Wen Ng, David W. Piston, Shankar Mukherji

## Abstract

The compartmentalization of eukaryotic cells into membrane-bound organelles with specific subcellular positioning enables precise spatial and temporal control of cellular functions. While functionally significant mitochondrial localization has been demonstrated in cells such as neurons, it remains unclear how general these cell principles are. Here, we examine the spatial organization of mitochondria within MIN6 pancreatic beta cells under variable glucose conditions. We observe glucose-dependent redistributions of mitochondria, favoring peripheral localization at elevated glucose levels when insulin secretion is also elevated. Our results, formalized into a stochastic model of mitochondrial trafficking, suggest that active mitochondrial transport along microtubules and calcium activity, but not ATP synthesis, are critical regulators of this redistribution. These results suggest that mitochondrial positioning may contribute to optimizing energy delivery in response to local demand, potentially representing a general regulatory mechanism across various cell types.

## Introduction

The organization of eukaryotic cells into organelles allows for spatiotemporal organization of biological processes, as seen via the specialized functionality of each respective organelle. Organelles are distributed throughout the cell in membrane-bound compartments and allow the cell to spatially segregate specific biochemical processes. Mitochondria, among the most dynamic organelles, are primarily responsible for generating energy in most eukaryotic cells. Elucidating how the biophysical context of mitochondria, in terms of both the morphology of the mitochondria itself as well as its interactions with the rest of the cell, regulate its cell biological functions remain frontier areas of investigation. Prior studies have mostly focused on mitochondrial morphology. Mitochondria behave like a constantly evolving network within the cell: the network can expand by undergoing fusion to other mitochondria and are broken down via fission or mitophagy in the case of damaged material^1-5^. The number, size, and genetic material (mtDNA) of individual mitochondrial structures have a well-established connection to the metabolic function of mitochondria and their ability to produce ATP^6-10^. These analyses, however, are more focused on properties of individual mitochondria with comparatively less emphasis on how they operate within the context of their host cells.

One potentially crucial, but less studied, factor coordinating mitochondrial activity and cellular function is the subcellular spatial positioning of mitochondria. Understanding how mitochondrial positioning responds to cellular demand may give us insight into how the cell optimizes organelle activity. Prior studies suggest that the cell appears to exert control over subcellular mitochondrial position. For example, it has been shown that cells express anchoring complexes that specifically exert control over mitochondrial positioning^11^. Furthermore, studies from cardiomyocytes have documented a preferred spatial allocation of mitochondria biomass within the cell^12^. Perhaps the most extensive observations of control over mitochondrial subcellular position have been shown in neurons, where both experimental and theoretical studies^13-16^ have indicated that neuronal mitochondria are trafficked to and retained in neuronal segments with high metabolic demand, typically in distal axons. The inference drawn from these studies is that since mitochondria produce energy for the cell, they preferentially position where those energy demands are highest. Given the highly polarized nature of neurons, whose elongated and highly branched geometry would significantly inhibit the cell’s ability to deliver adequate amounts of ATP by diffusion alone to distal axonal processes from the cell body, the need for active transport of mitochondria has a strong biophysical rationale. Whether this same logic can be applied to other, less polarized cell types is less examined and less clear, particularly in contexts where the energy demands are heterogenous even within less morphologically complex cells.

One such cell type that exhibits both relative morphological simplicity and strong spatial heterogeneity in energy demand is the pancreatic beta cell. In particular, glucose stimulated insulin secretion (GSIS) exhibits significant heterogeneity as the positioning of insulin granules are not random throughout the cell^17^. GSIS begins with the import of glucose into beta cells which leads to production of ATP that largely depends on the mitochondria^18^. The resulting change in ATP/ADP ratio drives the closing of K_ATP_ channels, which leads to membrane depolarization and calcium influx^19^. This influx of calcium is the ultimate step in driving insulin granule exocytosis. Because both closing K^+^ channels and the process of exocytosis require ATP at the cell periphery, there are reasons for the cell to preferentially localize its mitochondria there as well.

Here, we show that MIN6 cells show a subtle but significant shift in the spatial distribution of their mitochondria depending on their local glucose environment. Our results suggest that organelle positioning is an important factor of mitochondrial functionality, not only creating energy but also placing that energy where it is needed.

## Results

To assess the response of mitochondrial positioning to variations in functional demand on MIN6 cells, we established a quantitative framework for measuring mitochondrial location within the cell. To visualize the mitochondria, we incubated cells with MitoTracker CMX-ROS and carried out live-cell spinning disc confocal microscopy (Fig. 1A). We manually identified cell and nuclear boundaries (Fig. 1B,C), and then pre-processed the images using a combination of Gaussian and Laplacian filters to remove high frequency noise in the image. The denoised images were thresholded to segment mitochondrial pixels from the cellular background (Fig. 1D; Methods); we verified that our results are relatively insensitive to the detailed choice of thresholding and Gaussian blur parameters (Fig. S1). Crucially, to avoid having to decide exactly how to define mitochondrial position with a summary statistic derived from the segmented pixels, such as the position of the centroid of a particular mitochondrion, we instead measured the distribution of distances of all identified single mitochondrial pixels from the center of the nucleus (Fig. 1E,F). A pixel-level description of mitochondrial position allows us to focus on subcellular mitochondrial positioning independent of any potentially confounding effects from mitochondrial fission or fusion, which would not directly influence the location of mitochondrial pixels but would alter the calculated centroid position of individual mitochondria. For each cell, we normalized the distance of a given mitochondrial pixel by the maximum distance recorded between any mitochondrial pixel and the nucleus in that particular cell; we note that the mitochondrial pixel that displayed the maximum distance in each cell coincided with the cell periphery. The 1D coordinate system set up this way mitigates cell-to-cell variability in factors such as cell size, nucleus size, total mitochondrial mass, and the random of adherence to a glass-bottom dish. The relative position of a given mitochondrial pixel is thus also comparable from one cell to another: each pixel has a normalized distance value of at most 1.

**Figure 1.**
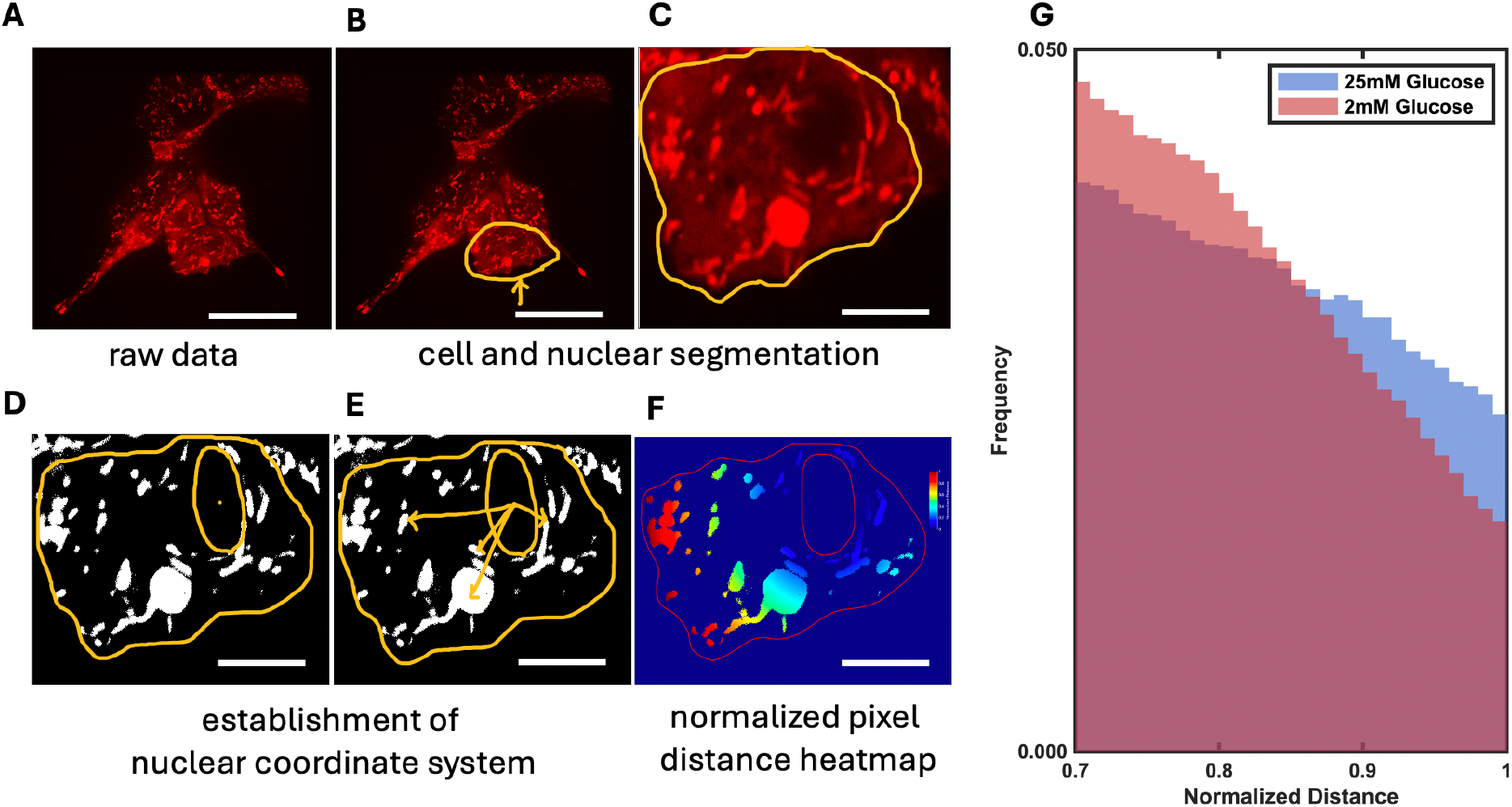
Mitochondrial spatial distribution in MIN6 cells under high and low glucose stimulation. Sample micrographs and schematic image analysis strategy of MitoTracker CMX-ROS labeled MIN6 cells: A) in the first subpanel we show a sample image of our raw image data, B,C) an individual cell is identified (yellow border), D) the nucleus is manually identified as a region of low fluorescence intensity, E) distances from the center of the nucleus to various mitochondrial pixels are indicated in the schematic, and F) we display a heatmap depicting the distances of each mitochondrial pixel to the center of the nucleus. In panels A and B, scalebar represents 20μm. In panels C-F, scalebar represents 10μm. G) Distribution of mitochondrial pixel distances from the center of the nucleus for the 30% most distant pixels from cells stimulated with 25mM (red histogram; N = 85 cells) and 2mM (blue histogram, N = 66 cells) glucose.

With this analysis, we were able to visualize pixel-level histograms of how nuclear-proximal versus membrane-proximal mitochondrial material is organized in space. We repeated these measurements in cells exposed to either low (2mM) or high (25mM) glucose concentrations. While the bulk of the pixel distributions do not show large differences between the two histograms, we observe a striking difference in the distributions at the peripheral edge of the cells (Figure 1G; Methods). Specifically, in 25mM glucose, MIN6 cells show a preference for peripheral mitochondria when exposed to 25mM glucose compared to 2mM glucose (Fig 1G). We note no significant differences in average number of mitochondria per cell or average mitochondrial size per cell (Fig. S2, Methods). The correlation of increased density of peripheral mitochondrial pixels with glucose, given that insulin secretion at the edge of the cell rises with glucose levels, is consistent with the hypothesis that mitochondria are actively positioned to meet local energy demands.

Having detected a difference between the levels of peripheral mitochondria as a function of glucose, we sought to uncover some of the mechanistic underpinnings of this pattern. Inspired by our hypothesis that mitochondria are positioned to serve as local energy supplies at sites of high energy demand, we tested three distinct factors that could drive spatial overlap between mitochondrial position and sites of demand: the generation of ATP in mitochondria, the increase of intracellular cAMP levels resulting from glucose stimulation, and the presence of a microtubule network that would bring mitochondria to the cell periphery.

To test the effects of ablating mitochondrial ATP production on mitochondrial position, we downregulated mitochondrial ATP production by treating cells cultured in 25mM glucose to the uncoupling agent FCCP. Interestingly, while we observe significant changes to mitochondrial morphology upon addition of 10nM FCCP (Fig. 2A), the fraction of mitochondrial pixels proximal to the edge of the cell does not change appreciably compared to when no FCCP is added, especially on the scale of the difference when varying glucose (Fig. 2B), nor do average mitochondria or number of mitochondria per cell (Fig. S3)

**Figure 2.**
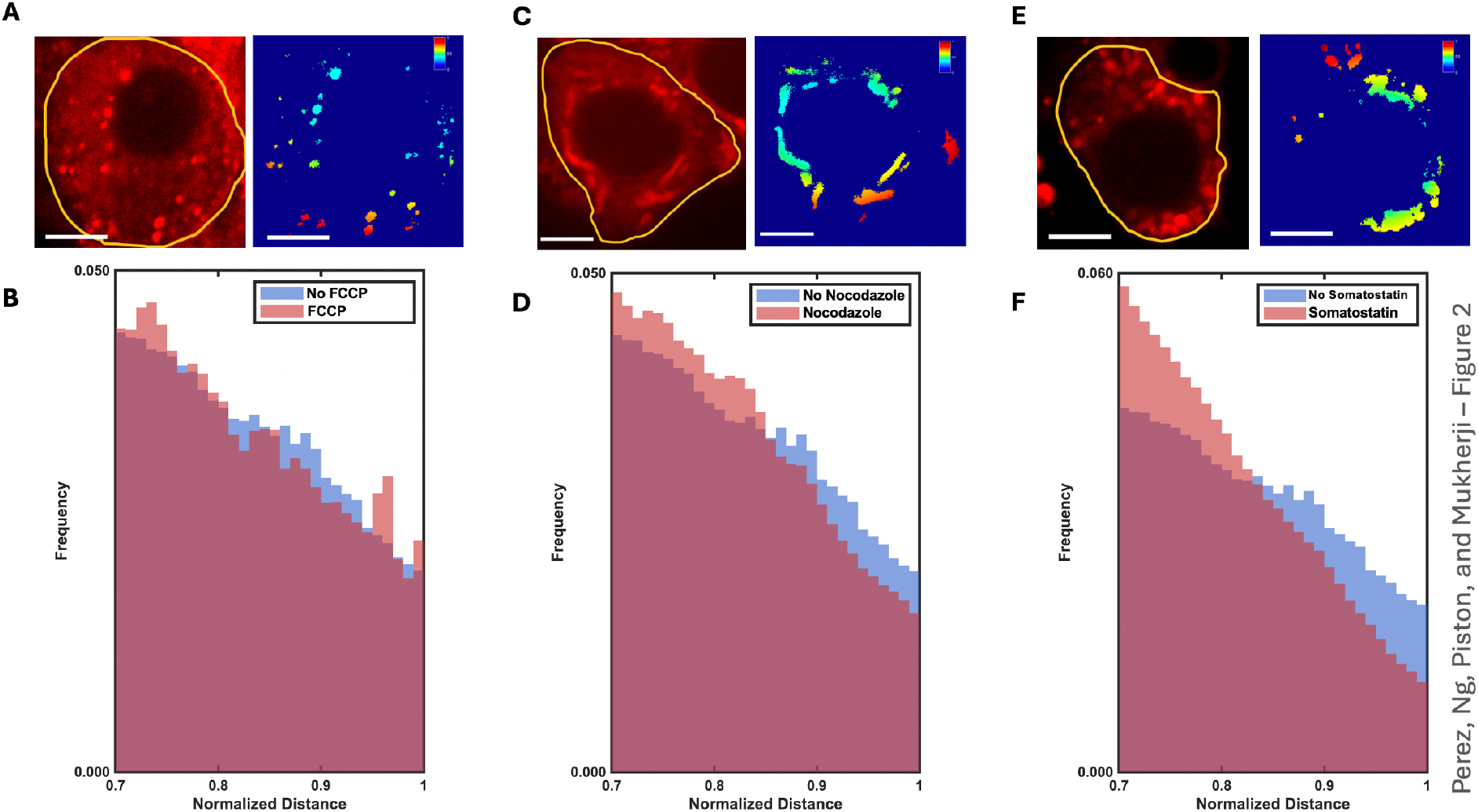
Responses of peripheral mitochondrial pixel distance distributions to perturbations in potential regulators of mitochondrial positioning. A) Example micrograph of mitochondrial morphology in response to treatment of MIN6 cells with 10nM FCCP. Scale bar represents 10μm. B) Response of mitochondrial pixel distribution to 10nM FCCP (blue histogram, N = 74 cells) compared to distribution observed in the absence of FCCP (red histogram repeated from Fig. 1B). C) Example micrograph of mitochondrial morphology in response to treatment of MIN6 cells with 10μM somatostatin. Scale bar represents 10μm. D) Response of mitochondrial pixel distribution to 10μM somatostatin (blue histogram, N = 72 cells) compared to distribution observed in the absence of somatostatin (red histogram repeated from Fig. 1B). E) Example micrograph of mitochondrial morphology in response to treatment of MIN6 cells with 10μM nocodazole. Scale bar represents 10μm. F) Response of mitochondrial pixel distribution to 10μM nocodazole (blue histogram, N = 62 cells) compared to distribution observed in the absence of nocodazole (red histogram repeated from Fig. 1B).

Next, to establish a role for active traffic in establishing glucose-stimulated trafficking of mitochondria to the cellular periphery^23,24^, we examined the mitochondrial pixel distribution in cells whose microtubule networks were disrupted. To effect this disruption, we treated the cells with 10 µM nocodazole, a microtubule destabilizer. As with other treatments, we observe no gross changes in mitochondrial morphology (Fig. 2C), or average mitochondrial number or size per cell (Fig. S4). With the microtubules depolymerized the histograms show a qualitatively similar response to treatment with. First, we observe a depletion in this peripheral region similar to our observation in low glucose (Fig. 2F). Second, however, we detect a sharp peak of mitochondrial at the cell periphery itself when compared to the control group of cells (Fig. 2D). This peak suggests a possibility that in the absence of the MT network, mitochondria retain an affinity for the cellular periphery that may play a role in the glucose-dependent shift of the mitochondrial pixel distribution.

Finally, to investigate mechanism underlying the signaling process that drives asymmetric mitochondrial traffic along the MT network toward the cell periphery as a function of glucose, we targeted the cAMP signaling pathway, a key downstream effector of mitochondrial ATP production^19-21^, on mitochondrial positioning. Glucose, for example, has been shown to drive an increase in cAMP levels, which in turn increase kinesin motor protein association and thus could play a role in glucose-dependent mitochondrial positioning. To this end we incubated our cells with the paracrine inhibitor somatostatin^22^. Somatostatin activates an inhibitory G-protein coupled receptor on beta cells and reduces insulin secretion, at least in part by reducing cAMP levels. Mitochondrial morphology as evaluated by visual inspection (Fig. 2E), and average number and size of mitochondria per cell (Fig. S5) does not appreciably change between cells treated and untreated with somatostatin on the timescale we measure. The mitochondrial histograms in response to 10uM somatostatin reveal a significant decrease in the density of mitochondrial pixels at the most peripheral region of the cells even in the presence of 25mM glucose, consistent with this reduced secretory demand (Fig. 2F).

Our experimental results suggest a minimal model in which mitochondrial interactions with the MT network and the cAMP signal transduction pathway play a crucial role in establishing mitochondrial positioning within the cell. To describe the mechanisms governing this dynamic localization, we developed a stochastic computational model to simulate mitochondrial movement and test whether it could qualitatively recreate the observed peripheral accumulation.

This model (Fig. 3A) explores the dynamic distribution of mitochondria among three states—free in the cytoplasm, bound to the cell membrane, or bound to microtubules—based on their binding and unbinding kinetics. The model operates in a one-dimensional spatial domain whose coordinate represents the percentage of the distance between the center and periphery of the modeled cell, thus simplifying the cell’s geometry to focus on central-to-peripheral movement. This 1D approach reduces computational complexity while capturing the core dynamics of mitochondrial movement. Mitochondria are initially distributed normally around x=50 (mean=50, std=10), reflecting their centralized positioning in low-glucose conditions. Free mitochondria undergo a random walk. Reflecting boundary conditions at x=0 and x=100 constrain mitochondrial positions within the domain, mimicking the physical barrier of the cell membrane.

**Figure 3.**
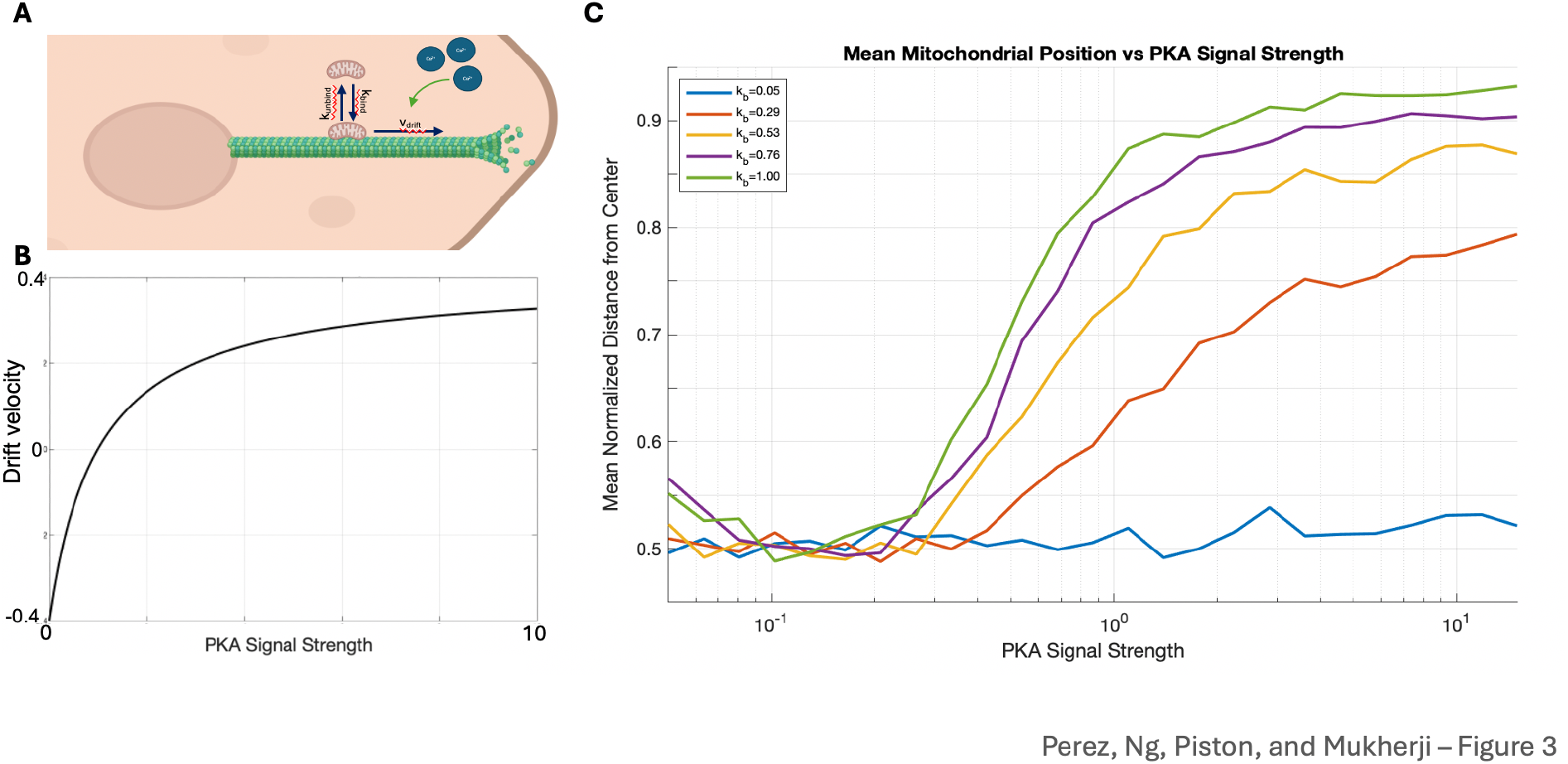
A) Schematic depicting mathematical model of mitochondrial interactions with cell periphery and microtubules, with a third cytoplasmic state in which mitochondria are localized to neither the membrane or microtubules. B, C) Steady state results of mitochondrial binding to target compartments: Fraction of mitochondrial localized to cell periphery/membrane (B) or microtubules (C) as a function of the ratio of mitochondrial affinities for the membrane (α) and for the microtubules (β). D) Slice from panels B and C in which the affinity for the membrane is fixed and the affinity for the microtubules is varied; note that at 0 affinity for the microtubules we observe a maximum in the fraction of mitochondria at the membrane, but with the fraction less than 1.

The model’s core feature is its representation of cAMP/PKA/Ca2+-dependent transport along MTs, designed to capture the glucose-induced peripheral trafficking observed in MIN6 cells (Fig. 3A). Mitochondria transition stochastically between free and MT-bound states. The binding rate is spatially variable to account for higher MT density near the cell center. With these kinetics governing the mitochondrial motion, we can define the drift velocity of MT-bound mitochondria as:

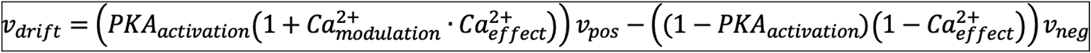

To examine the model, we varied the cAMP/PKA_signal_ over 25 logarithmically spaced values from 0.05 to 15, mapping to PKA_activation_ via a Hill function. The Ca^2+^ signal spans 5 linearly spaced values from 0.1 to 2.0, which enhances kinesin activity and inhibits dynein (Fig. 3A). Specifically, we observe that in the model high PKA_signal_ and Ca^2+^ signal increase v_drift_, which thereby breaks the symmetry of the mitochondrial random walk and results in driving mitochondria to the periphery (Fig. 3B).

We explored a comprehensive parameter grid with k_binding, constant_ having 25 values from 0.05 to 1.0, PKA_signal_ similarly having 25 values from 0.05 to 15) and Ca _signal2+_ with 5 values, 0.1 to 2.0. These values were selected to cover a broad parameters space, yielding 3125 combinations to cover a wide range of glucose and microtubule conditions. In order to compare the model output to our experimental results, we computed the mean mitochondrial position over each set of simulations for given parameter values and plotted these mean positions as a function of PKA_signal_ and k_binding_. Overall, the model recapitulates both the MT- and calcium signaling-dependent shift of mitochondrial density to the periphery we observed experimentally (Fig. 3C). At very low binding (Fig. 3C, blue curve) we observe that the mitochondrial position remains near the middle of the cell no matter the PKA_signal_ value, while for appreciable binding of mitochondria for microtubules we observe an increase in peripheral mitochondria that scales with the strength of PKA_signal_, thus describing the dependence of mitochondrial positioning on both the microtubule network as well as Ca^2+^/cAMP/PKA signaling.

## Discussion

Mitochondria produce ATP through oxidative phosphorylation and careful work has drawn links between mitochondrial-specific geometric properties and the rate with which mitochondria generate ATP^25,26^. Polarized structures, such as neurons, muscle cells, and heart cells are known to have mitochondria spatial distribution correlated with regions of high energy demand. These cells share the common feature of signal transduction, with a clear need for temporal and spatial efficiency in order to communicate with neighboring cells. Whether these same principles play out in cell types whose morphologies are comparatively less polarized is unclear. We sought to use GSIS in pancreatic beta cell-like MIN6 cells as a model system to explore the response of mitochondrial position to elevated energy demand.

We observed that in conditions of high secretory, and thus energy, demand there was a significant difference in the mitochondrial density at the furthest edges of these cells. This shift in mitochondrial density appears to be primarily dependent on glucose and the presence of an intact MT network. We were able to qualitatively recapitulate our experimental patterns in a mathematical model that described a competition for mitochondria between the MT and the cell periphery whose affinity for mitochondria is glucose-dependent, which can augment previous efforts to capture mitochondrial ATP synthesis. This retention may help ensure mitochondrial energy production supports insulin secretion, as localized energy may be necessary for efficient exocytosis of insulin granules. We note also that mitochondrial function extends beyond ATP production; in particular, mitochondria are major regulators of intracellular calcium levels, which play a major role in insulin secretion^19-21^ and our somatostatin results suggest that cAMP levels, and thus calcium, could also help regulate mitochondrial position. While we have not uncovered the mechanism by which this glucose-dependent affinity for the cell periphery is effected, we believe that our quantitative analysis strategy will aid future searches for this putative regulator of mitochondrial positioning.

In the broader context of the interplay between mitochondria and insulin secretion, we note that insulin granules exhibit similar dynamics to what we infer mitochondria obey from our analysis. It was recently shown, for example, that destabilizing the microtubule network via nocodazole treatment inhibits the rate at which insulin granules are withdrawn from the cell periphery^27^, matching the pattern we observe in mitochondria, which exhibit a spike in density at the cell periphery upon nocodazole treatment. Indeed, numerous studies have shown that disruption of the actin cytoskeleton, both in the bulk of the cell but especially with disassembly of cortical actin, enhances GSIS^28-31^. While these results have been interpreted as largely a consequence of increased access for insulin granules to sites of exocytosis, it is possible that the mitochondrial peripheral positioning we observe also plays a role. In the language of our mathematical model, the glucose-dependent MT instability could tilt the competition for mitochondria away from the MT and thus bulk of the cell and toward the membrane. This picture is consistent with the correlation we observe between mitochondrial peripheral positioning to conditions that result in higher insulin secretion.

From the perspective of the cell and its mitochondria, one potential benefit to a competitive-binding based mechanism of mitochondrial positioning is the flexibility it affords to the cell in terms of allowing individual mitochondria within the cell to perform the wide variety of biochemical tasks assigned to them. Elevated mitochondrial fusion, for example, globally links mitochondria together into a networked structure well suited to supplying the entirely of the cell with ATP during periods of elevated cellular proliferation^32^, but potentially at the cost of homogenizing the mitochondrial compartment, rendering it unable to perform otherwise incompatible biochemical processes simultaneously^10^. MIN6 cells, and their pancreatic beta cell counterparts, however, need to balance the calcium and energetic maintenance functions that mitochondria provide to the cell with the increased demands placed on them by glucose stimulation. Competitive-binding based mitochondrial positioning in cells such as MIN6 and their pancreatic beta cell counterparts, however, could allow the cell to not be required to extensively remodel mitochondrial composition to elevate ATP production rates needed to boost insulin secretion, but simply capture them at sites of increased demand to elevate local ATP concentrations.

Finally, while our results and analysis have been focused on MIN6 cells, we note that the phenomenon of spatially inhomogeneous energy demands on cells upon external or internal cues is a widespread phenomenon. From the highly demanding secretory activity of B cells to the epithelia lining the gut, we suggest that mitochondrial positioning may play an important role in matching local energy supply with energy demand in a wide variety of cell biological contexts.

## Acknowledgements

We thank A. Ustione for assistance with microscopy and members of the Piston and Mukherji groups for discussions and critical evaluation of the manuscript. This work was supported by the Washington University Imaging, Modeling and Engineering of Diabetic Tissues Training Grant T32DK108742 (to L.P.), R01DK123301 (to D.W.P.), and R35GM142704 (to S.M.).

## METHODS

*Cell Culture* MIN6 cells were grown in Dulbecco’s modified Eagle’s medium (DMEM, 25 mmol/l glucose) equilibrated with 5% CO2 and 95% air at 37 C. The medium was supplemented with 15% fetal bovine serum (FBS) and 5% penicillin and streptomycin. MIN6 cells used in this study were harvested at passages 30-40.

*Analysis of Mitochondrial Properties* Semiconfluent cells were grown on a separate dish in anticipation for imaging. On the day of the experiment, cells in their respective dishes were incubated with 100 nanomolar MitoTracker Red CMXRos for one hour prior to imaging, adding the dye to a fresh aliquot of regular media the cells were grown in. After one hour, the cells were rinsed with KRBH, which behaved as a clear imaging media for the cells.

*Imaging Protocol* Cells were imaged using a Nikon Spinning Disc Confocal Microscope, fully equipped with an incubator (Tokai-Hit) enabling full temperature, CO2, and humidity control. Cells were grown in glass bottom dishes, which was fixed on a microscope stage, and images were obtained using a PlanApo 100x 1.45 NA oil-immersion objective. Z stacks were obtained with a z spacing of 0.2 um.

The cells were maintained at 37 degrees Celcius and 5% CO2. The Red CMXRos dye was excited with a 561nM laser and fluorescence emission was detected through a 665nM filter (red emission channel). Unless otherwise stated, gain levels, confocal aperture size, and laser power were adjusted to not saturate the detector, but remained constant throughout the time-series of image acquisition.

*Image Analysis* Images were opened up using Fiji, where the entire field of view of cells was visible. We identified a cell that only minimally overlapped with a neighboring cell, and then cropped this cell. This cropped cell was analyzed using a custom MATLAB script that identifies all mitochondrial pixels within the cell. Mitochondrial pixel identification begins with applying a 2D Gaussian smoothing filter to each Z slice of the fluorescence image data to denoise the image data (using a smoothing radius σ = 2.5) followed by a Laplacian filter to sharpen the image. Following this pre-processing step, we create a binary mask of pixels whose fluorescence value is above a fixed threshold value (750 for our setup) for each Z slice of the fluorescence image. We then use the binary mask to identify mitochondria as 3D connected components defined using only pixels whose 26 neighboring pixels in 3D are also included in the mask using the MATLAB bwconncomp command. We then filter the connected components to remove components that are too small (125 pixels or less) or too large (100000 pixels or more) to represent mitochondria. We define the number of mitochondria in a cell to be the number of connected components in the cell obtained from the filtered connected component analysis, and the average size of the mitochondria in a cell to be the average number of pixels across the connected components within a cell. These results are reported in Supplementary Figures S2-5. The mitochondrial pixels used in the distance analysis belongs to these filtered connected components identified from the fluorescence images.

To carry out the distance measurements, we begin by manually setting the center of the nucleus as the origin for each respective cell. For a given cell, for each identified mitochondrial pixel, we compute the Euclidean distance to the center of the nucleus. We divide this Euclidean distance by the maximum pixel distance in the given cell under analysis to obtain the normalized distance. The set of normalized distances are then used to construct the histograms shown in Figures 1 and 2.

*Titration Experiments* On day prior to the experiment, MIN6 cells from a T-25 flask were monitored to verify the approximate optimal confluency for the cell line at 70-80% density. We incubated the cells with Trypsin-EDTA (1x) for 15 minutes, or until the cell layer was dispersed, to allow for proper aliquoting, with a similar optimal density profile, to culture vessels suitable for imaging. For static incubation, the cells were incubated in their normal media as outlined above. On the day of the experiment, cells were preincubated in the respective perturbation conditions- 2mM Glucose, 10nM FCCP, 10μM somatostatin, and 10μM nocodazole- for 60 minutes. Just prior to imaging, culture medium was rinsed and replace with the appropriate imaging media (1x KRBH). This was supplemented with the respective conditions of the titration to ensure maintenance of cellular environment during live imaging. Cells in the dish were then placed in the incubator and underwent live-cell imaging in 5% CO2 conditions at 37C.

