## Supplementary Figures for "Mitochondrial position responds to glucose stimulation in a model of the pancreatic beta cell"

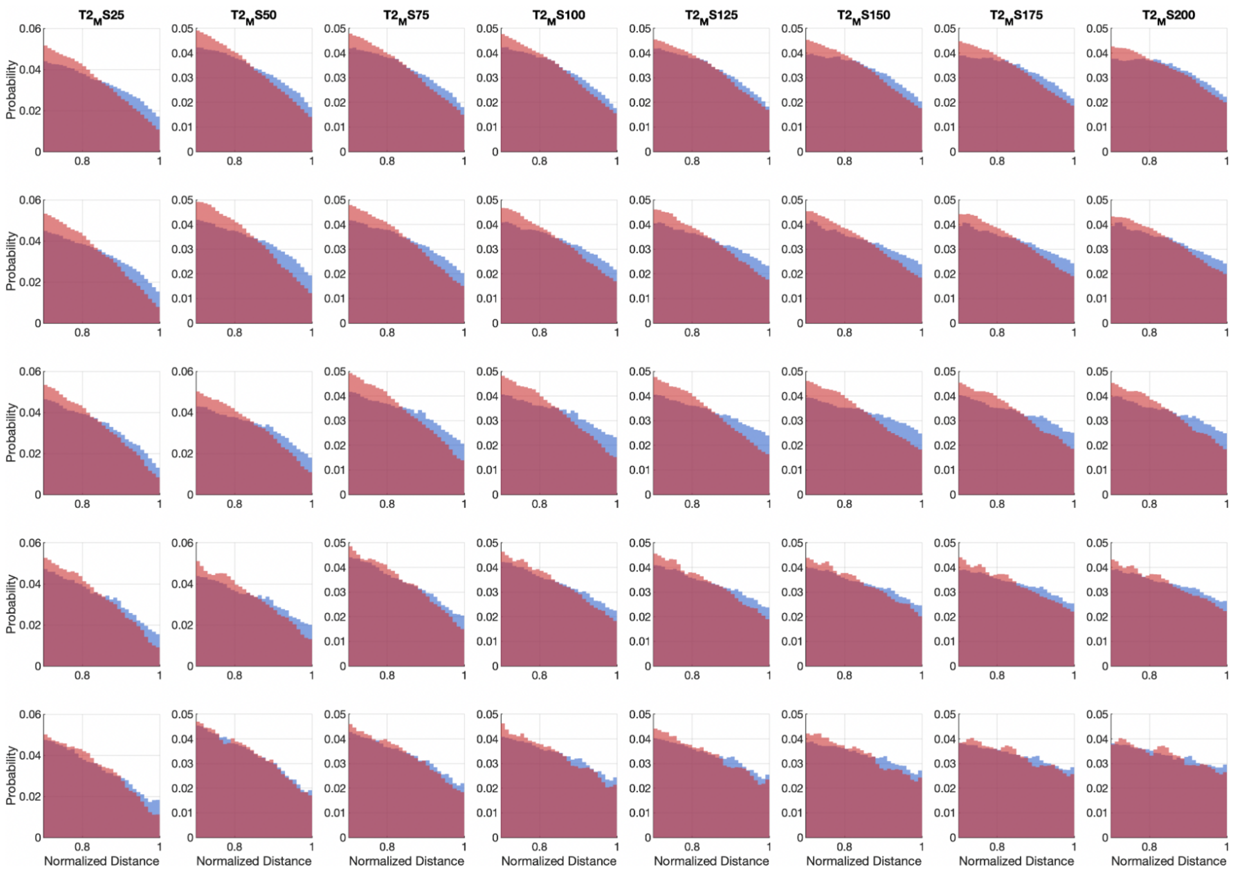


Figure S1. Sensitivity analysis of mitochondrial pixel probability distributions as a function of smoothing and thresholding parameters. Connected component size thresholding parameters are varied across columns between 25 to 200 pixels, fluorescence intensity thresholding parameters are varied across rows from 100-1000.

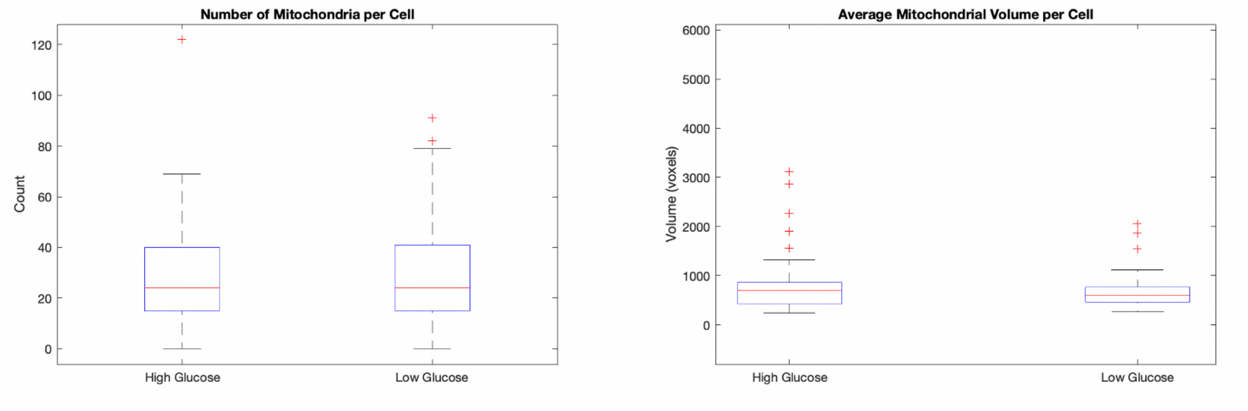


Figure S2. Box and whisker plots showing the median (red line) and 50% interquartile range (blue boxes) of number of mitochondria per cell and average mitochondrial volume per cell for cells cultured in 25mM glucose (high glucose) and 2mM glucose (low glucose)

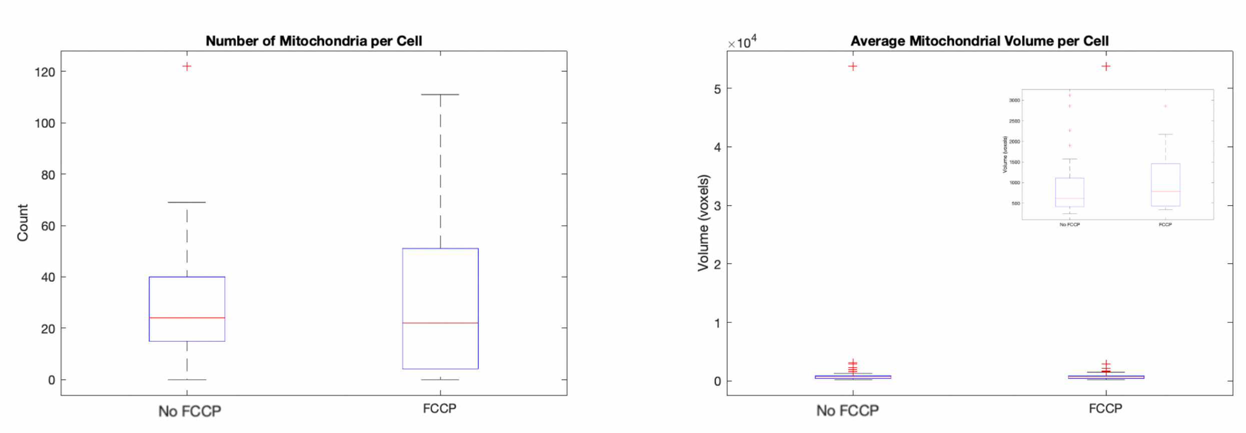


Figure S3. Box and whisker plots showing the median (red line) and 50% interquartile range (blue boxes) of number of mitochondria per cell and average mitochondrial volume per cell in cells cultured in the absence and presence of the uncoupling agent FCCP. Inset in right panel shows box and whisker plots with points with outlier average mitochondrial volume datapoints removed to facilitate comparison between the two populations.

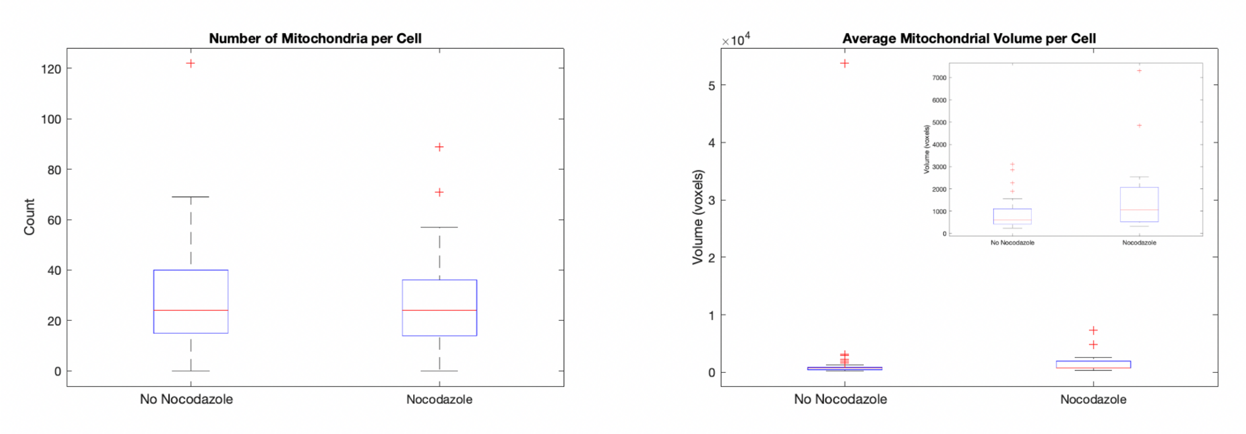


Figure S4. Box and whisker plots showing the median (red line) and 50% interquartile range (blue boxes) of number of mitochondria per cell and average mitochondrial volume per cell in cells cultured in the absence and presence of the microtubule disrupting agent nocodazole. Inset in right panel shows box and whisker plots with points with outlier average mitochondrial volume datapoints removed to facilitate comparison between the two populations.

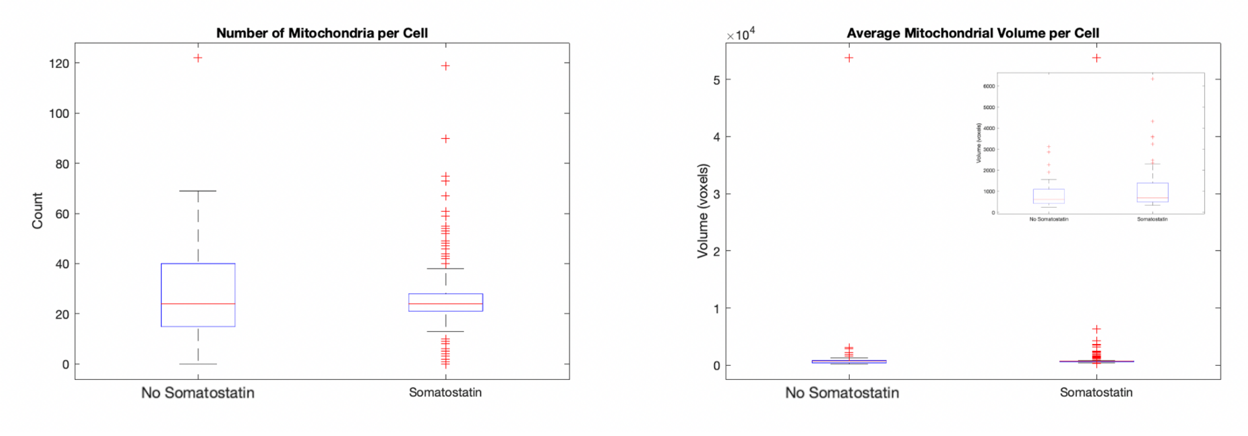


Figure S5. Box and whisker plots showing the median (red line) and 50% interquartile range (blue boxes) of number of mitochondria per cell and average mitochondrial volume per cell in cells cultured in the absence and presence of somatostatin. Inset in right panel shows box and whisker plots with points with outlier average mitochondrial volume datapoints removed to facilitate comparison between the two populations.
